# Acute stress yields a sex-dependent facilitation of signaled active avoidance in rats

**DOI:** 10.1101/2024.04.27.591470

**Authors:** Samantha L. Plas, Cecily R. Oleksiak, Claire Pitre, Chance Melton, Justin M. Moscarello, Stephen Maren

## Abstract

Post-traumatic stress disorder (PTSD) is a debilitating disorder characterized by excessive fear, hypervigilance, and avoidance of thoughts, situations or reminders of the trauma. Among these symptoms, relatively little is known about the etiology of pathological avoidance. Here we sought to determine whether acute stress influences avoidant behavior in adult male and female rats. We used a stress procedure (unsignaled footshock) that is known to induce long-term sensitization of fear and potentiate aversive learning. Rats were submitted to the stress procedure and, one week later, underwent two-way signaled active avoidance conditioning (SAA). In this task, rats learn to prevent an aversive outcome (shock) by performing a shuttling response when exposed to a warning signal (tone). We found that acute stress significantly enhanced SAA acquisition rate in females, but not males. Female rats exhibited significantly greater avoidance responding on the first day of training relative to controls, reaching similar levels of performance by the second day. Males that underwent the stress procedure showed similar rates of acquisition to controls but exhibited resistance to extinction. This was manifest as both elevated avoidance and intertrial responding across extinction days relative to non-stressed controls, an effect that was not observed in females. In a second experiment, acute stress sensitized footshock unconditioned responses in males, not females. However, males and females exhibited similar levels of stress-enhanced fear learning (SEFL), which was expressed as sensitized freezing to a shock-paired context. Together, these results reveal that acute stress facilitates SAA performance in both male and female rats, though the nature of this effect is different in the two sexes. We did not observe sex differences in SEFL, suggesting that the stress-induced sex difference in performance was selective for instrumental avoidance. Future work will elucidate the neurobiological mechanisms underlying the differential effect of stress on instrumental avoidance in male and female rats.

## INTRODUCTION

Extreme stress associated with witnessing or experiencing a life-threatening event can lead to post-traumatic stress disorder (PTSD), which is nearly twice as common in women compared to men (Olff, 2017). Symptoms of PTSD include hyperarousal, intrusive memories and flashbacks, alterations in mood and cognition, and pathological avoidance behavior (5th ed.; DSM–5; 2013). Among these symptoms, avoidance is particularly problematic because it limits an individual’s opportunity to experience trauma reminders in a safe setting. As a result, avoidance behavior maintains PTSD symptoms by preventing the extinction of traumatic fear memories (Thwaites and Freeston, 2005). Although avoidance plays a critical role in the etiology of PTSD, little is known about how traumatic stress contributes to the emergence of avoidance behavior.

In the laboratory, avoidance can be modeled in animals using avoidance learning procedures. In active avoidance tasks, animals must perform a specific behavioral response to avoid danger, whereas in passive avoidance tasks animals must withhold a behavioral response that leads to an aversive outcome (Krypotos et al., 2015; Diehl et al., 2019; Lopez-Maraga et al., 2022). Some of the earliest work examining the relationship between avoidance and stress demonstrated that experience with an acute and inescapable footshock (a stressor) undermined subsequent escape and avoidance responding (Brown and Jacobs, 1949; Carlson and Black, 1960; Leaf, 1964), an effect that was later attributed to “learned helplessness” (Overmier and Seligman, 1967). However, subsequent work in rats revealed conflicting effects of stress on avoidance learning. In some studies, inescapable shock was found to impair avoidance learning (Seligman et al., 1975; Weiss and Glazer, 1975), whereas in others it facilitated avoidance learning (Koba et al., 2001; Brennan et al., 2005). The conflicting nature of studies examining the impact of stress on avoidance is likely due to difference in avoidance procedures (active vs passive), type of stressor (acute vs chronic), and the model system being used (For review see Lopez-Moraga et al., 2022). Furthermore, many of these studies failed to consider sex as a biological variable. This is particularly problematic given that women engage in avoidant coping mechanisms more than men (Kessler et al., 1995; Fullerton et al., 2001; Sheynin et al., 2017; Panayiotou et al., 2017). Unfortunately, studies that have used both male and female subjects also yield conflicting results (Dalla et al., 2007; Shors et al., 2007, McDowell et al., 2015, Collins et al., 2023).

Here, we aimed to resolve this matter by assessing the impact of inescapable shock (an acute stress procedure) on the acquisition and extinction of two-way signaled active avoidance (SAA) in male and female rats. In SAA, rats learn to shuttle in response to a conditioned stimulus (CS, tone) to avoid a forthcoming aversive unconditioned stimulus (US, footshock). Although instrumental avoidance is adaptive, it is mediated by behavioral contingencies that are hypothesized to underlie pathological avoidance in humans with PTSD (Krypotos et al., 2015; Moscarello and Hartley, 2017; Ball and Gunaydin, 2022). For this reason, instrumental avoidance serves as an important model for understanding the behavioral and neural mechanisms by which stress and trauma lead to pathological avoidance behavior. To assess the impact of stress on SAA, we employed an acute stress procedure (i.e., inescapable footshock) that has previously been shown to sensitize aversive Pavlovian fear conditioning, a phenomenon termed stress-enhanced fear learning (SEFL). This stress procedure models several key features of PTSD including the sensitization of future fear learning and anxiety-like behaviors (Poulos et al., 2014; Rajbhandari et al., 2018; for review see Nishimura et al., 2022). Notably, the SEFL phenomenon is long-lasting. It persists for at least 90 days in adult rats (Rau & Fanselow, 2009) and through adulthood when delivered as a form of earlylife stress (Quinn et al., 2013; Poulos et al., 2014). Although the ability of traumatic stress to yield SEFL is well characterized, there is less known about how it influences instrumental avoidance learning, including SAA.

In two experiments we examined the influence of acute stress (inescapable shock) in the acquisition and extinction of instrumental avoidance (Experiment 1) and on SEFL using a Pavlovian fear conditioning procedure (Experiment 2) in male and female rats. From a clinical perspective, stress might be expected to facilitate avoidance responding, given that avoidance behavior is a core symptom of stressorand traumarelated disorders including PTSD. Alternatively, stress might undermine avoidance behavior by promoting competing behaviors, including freezing (Moscarello and Hartley. 2017; Livermore et al., 2021; Hoffman et al., 2022) or by inducing learned helplessness (Overmier and Seligman, 1967). Indeed, SEFL manifests as enhanced conditioned freezing in response to a single footshock (Poulos et al., 2015; Rajbhandari et al., 2018). The experiments reported here support the first possibility and reveal that acute stress facilitates the acquisition and performance of SAA in a sex-dependent manner despite producing similar levels of SEFL in male and female rats.

## MATERIALS AND METHODS

### Animals

The subjects were male (Experiment 1: 54; Experiment 2: 20; 200-250 g) and female (Experiment 1: 30; Experiment 2: 20; 200-250 g) Sprague-Dawley rats (Taconic Bioscience Inc.) roughly PN60 days of age upon arrival. All rats were individually housed in a temperature and humidity-controlled vivarium, with a 14h/10h light/dark cycle and ad libitum access to standard rodent chow and water. Behavioral testing was conducted during the light phases. Rats were handled for 2 min/d for 5 d before behavioral testing to acclimate them to the experimenter. All experiments were conducted in accordance with the National Institutes of Health guide for the care and use of Laboratory animals.

### Behavioral procedures and apparatus

**Experiment 1**. Male and female rats were assigned to either stress (n = 44) or no-stress conditions (n = 40). For the acute stress procedure, rats were transported from the vivarium, placed in the acute stress context, and given 15 unsignaled shocks (1 s, 1 mA), with an average inter-shock interval of 6 min distributed over a 90-min period. Shock delivery was controlled by Med-PC (Med Associates). Control animals were placed in this same context for 90 minutes without shock. The acute stress procedure was conducted in eight identical rodent conditioning chambers (30 cm × 24 cm × 21 cm; Med Associates) housed in sound-attenuating cabinets used to deliver inescapable shocks (i.e., the stress procedure). The chambers consisted of two aluminum side walls, Plexiglas ceilings and rear walls, and a hinged Plexiglas door. A 15-W white house light within each chamber was illuminated and the room was illuminated with white overhead fluorescent lights; the doors of the sound attenuating cabinets were closed. A grid floor consisting of 36 stainless steel rods was wired to a shock source and a solid-state grid scrambler (Med Associates) for delivery of footshock USs. Each chamber rested on a load cell force transducer which measured displacement of the chamber in response to each rat’s motor activity. Load cell voltages were acquired at 5 Hz, thus allowing for assessment of locomotor activity every 200 msec. These voltages were digitized and transformed to absolute values (0 – 100 units). Freezing was quantified by the number of observations below the freezing threshold (< 10). A freezing bout was defined as 5 consecutive values below this threshold, corresponding to 1 s of freezing.

One-week later rats underwent four days of SAA training in a standard shuttle box apparatus. The shuttle boxes (50.8 × 25.4 × 30.5 cm, l × w × h; Coulbourn Instruments) were constructed of Plexiglas and metal and were divided across their long axis into two even compartments by a metal divider with an open passage (8 × 9 cm, w × h) to allow animals to cross freely between two compartments. The floors consisted of conductive stainless-steel bars through which electric shock was delivered. Two speakers, integrated into the walls opposite the divider, delivered a 2-kHz, 80dB tone CS for 15 seconds. A scrambled footshock US (0.7-mA, 0.5-s) was delivered through the grid floor. Each shuttle box was contained inside a larger sound-attenuating chamber. The shuttle-box context used for SAA training featured no house light or compartment lights, and black construction paper wall inserts with glow-in-the-dark stars. This context was cleaned with 1% ammonium hydroxide, a small volume of which was placed in the waste pan under each grid floor. The doors of the sound attenuating chambers were left open. Rats were transported from the vivarium to the test room in black plastic bins without bedding. On the first day of training, rats were presented with a single Pavlovian CS-US trial; this ensured that the animals experienced at least one tone-shock pairing. Following this trial, and during subsequent conditioning sessions (days 2-4), training consisted of 30 avoidance trials in which an avoidance response (shuttling during the CS) terminated the CS and omitted the US. The intertrial interval during training was on average 120 seconds. Stimulus delivery was controlled by GraphicState Software (Coulbourn Instruments). Two infrared emitter-detector arrays, one in each compartment, were used to monitor shuttling (crossing from one compartment to the other in either direction). Coulbourn GraphicState software (Coulbourn Instruments) both controlled stimulus delivery and acquired shuttling responses.

After training was completed, male and female animals from stress and no-stress conditions were assigned to groups with matching avoidance performance before receiving extinction in either the SAA training context or a novel shuttle-box context. To distinguish it from the SAA training context, the novel shuttle-box context had an illuminated house light in each compartment and black and white striped wall inserts. The shuttle boxes were cleaned with 3 % acetic acid, a small volume of which was placed in the waste pan under each grid floor. A black Plexiglas floor insert was added over the grid. Rats were transported from the vivarium to the test room in white plastic bins with bedding. There were 3 daily sessions of extinction, during which 30 CSs were presented in the absence of the US. In addition, CSs did not terminate with shuttling during extinction.

**Experiment 2**. Male and female rats were assigned to either stress (n = 20) or no-stress conditions (n = 20). The acute stress procedure and control procedure were the same as above. A week after these procedures, rats underwent a one-trial contextual fear conditioning procedure in a different set of eight rodent conditioning chambers. To distinguish the context for the one-trial contextual fear conditioning procedure from the acute stress context (described above), black and white striped wall inserts were used, and the room was illuminated with red overhead fluorescent lights. A grid floor of 19 stainless steel rods was used and the doors of the sound attenuating cabinets were left open. The chambers were cleaned with 3% acetic acid solution, and a small volume of this solution was placed in the waste pan under each grid floor. Rats were transported from the vivarium to the test room in white plastic boxes without bedding. The one-trial contextual fear conditioning procedure involved one unsignaled shock (1 s, 1 mA) after a 3-minute baseline period. Subjects were then removed from the chamber 30 seconds after the shock. Twenty-four hours later, rats were returned to the same context for a fear retrieval test that lasted 8 minutes. Twenty-four hours later, rats were returned to the acute stress context for a fear retrieval test that lasted 8 minutes.

### Statistical analysis

SAA learning was quantified using the total number of avoidance responses (ARs) made during each training session. Rats were considered poor avoiders if they avoided an average of six or fewer times on the last three days of training. For all extinction tests, the number of ARs were capped at 1 response per CS for a total of 30 possible per test to match training. The number of intertrial responses (ITRs) were reported as a sum of responses during each trial. Performance was independent of the extinction context, so the groups were collapsed. Contextual fear conditioning was assessed by quantifying the percentage of time spent freezing during the pre-shock period (3 minutes) and post-shock period (30 seconds). For contextual fear retrieval tests, freezing was quantified and averaged across the 8-min test. ANOVAs followed by post hoc tests were performed on all data. All statistical analyses were performed in Statview version 5.0.1 (SAS institute) in a MacOS 9 open-source emulator.

## RESULTS

### Experiment 1: Experiment 1: Effect of acute stress on acquisition and extinction of SAA

Experiment 1 aimed to examine the role of acute stress on the acquisition and extinction of SAA. To this end, we submitted rats of both sexes to either an acute stress or control procedure. Rats then received four days of SAA training, beginning one week after acute stress (Figure 1A). As shown in Figure 2A and 2B, acute stress increased the acquisition of SAA in female but not male rats. A two-way ANOVA with a between-subjects factor of Treatment (Stress or Control) and a within-subjects factor of Day was performed on the data from each sex. In females, the ANOVA revealed a main effect of Treatment [F(1, 28) = 9.56, p = 0.0045] and Day [F(3, 84) = 49.44, p < 0.0001], indicating that stress significantly increases ARs in females across the training session. In contrast, stress did not affect performance in male rats [F(1, 52) = 0.327, p = 0.5701]. However, a main effect of Day [F(3, 156) = 52.06, p < 0.0001] revealed that male rats successfully acquired the avoidance response. Interestingly, there was also a sex difference in the proportion of rats that failed to acquire the avoidance response (“poor avoiders”). As shown in Figure 2A, there were no poor avoiders in females that underwent the stress procedure, compared to 2 of 14 rats in the control condition. In males (Figure 2B), however, there were similar proportions of poor avoiders in the Stress and Control groups. Thus, acute stress prevented the emergence of poor avoiders and facilitated acquisition of SAA in females, but not males.

**Figure 1.**
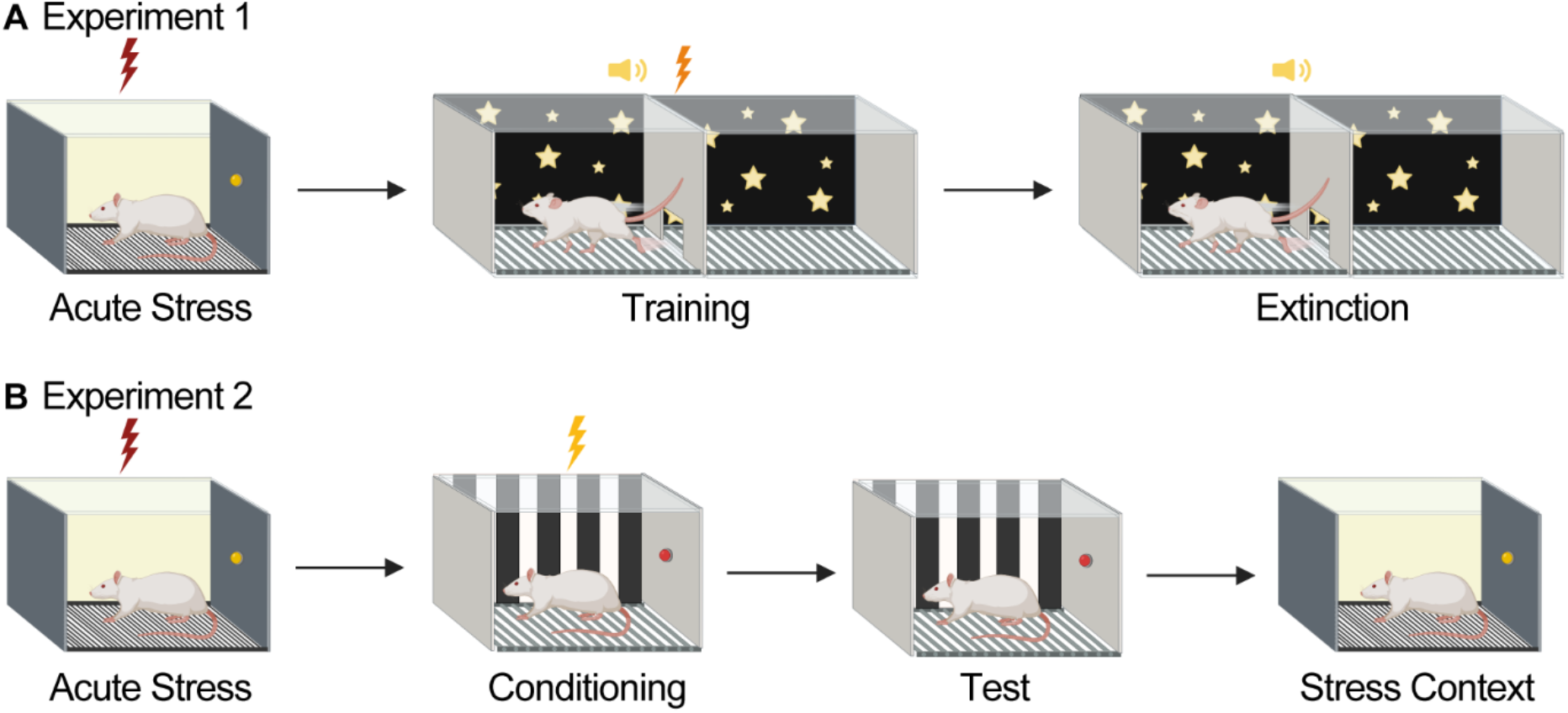
Behavioral Design. (A) Experiment 1. Rats were submitted to acute stress or control procedures (no shock). One week later, rats received 4 days of two-way signaled active avoidance (SAA) training. Subjects then received 3 days of SAA extinction in either the training context or a novel context (performance did not differ in the two extinction contexts, and data were collapsed). (B) Experiment 2. Rats were submitted to acute stress or control procedures. A week later, rats were submitted to a one-trial contextual fear conditioning paradigm in which 1 US was delivered. The following day, freezing was examined to test for the presence of stress enhanced fear learning (SEFL). 24 hours later rats were placed back into the stress context for a stress context retrieval test (created with Biorender.com).

**Figure 2.**
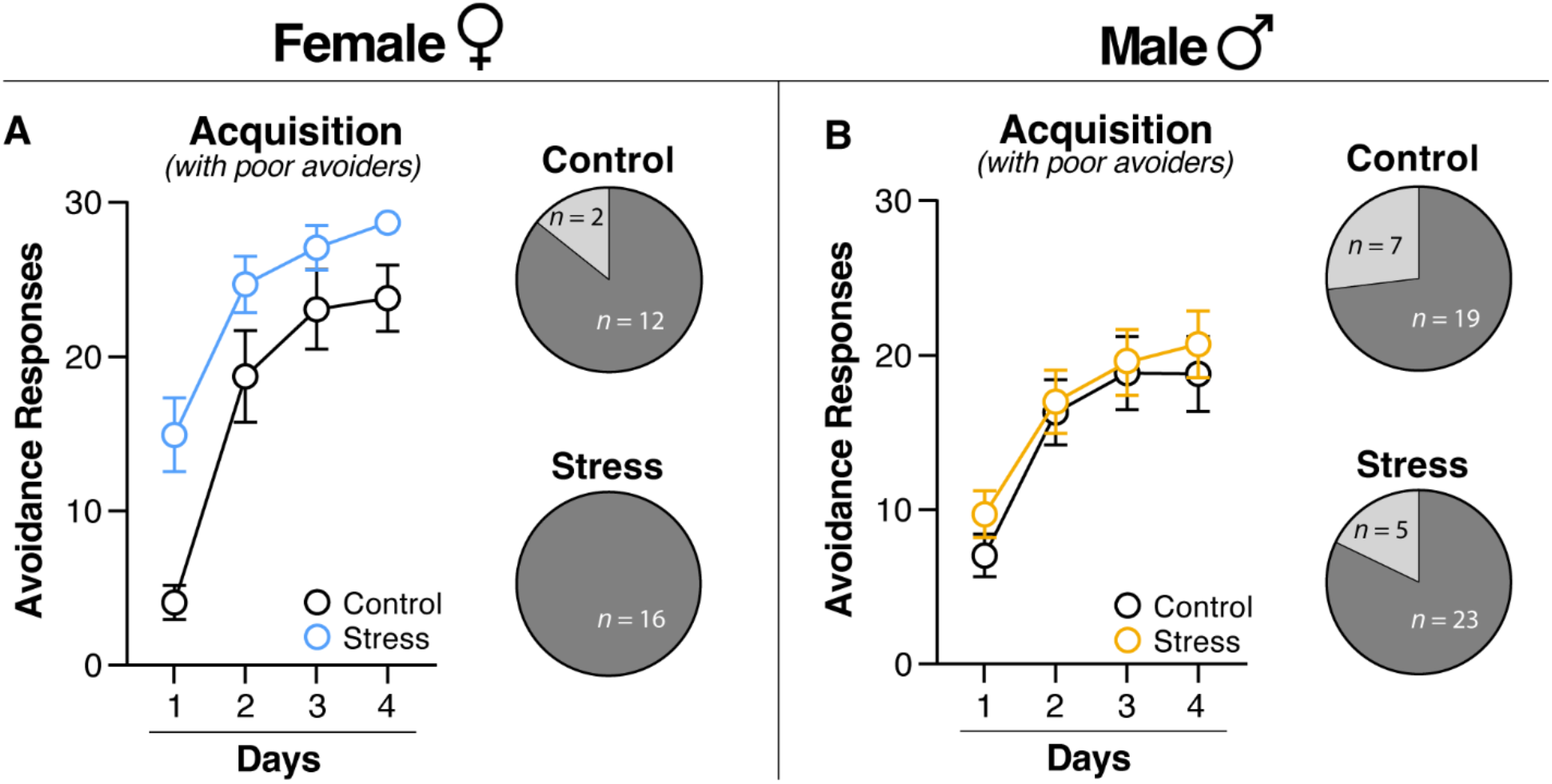
Acute stress facilitates acquisition of two-way signaled active avoidance in female rats. (A) Female and (B) Male (Left) average number of ARs performed on each day of SAA training including poor avoiders. (Right) Pie charts representing the number of good and poor avoiders per group. (A) Female rats exposed to acute stress performed significantly higher ARs than their control counterparts during SAA training. (B) Male rats submitted to acute stress performed similarly to their control counterparts during SAA training. As seen in the pie graphs, poor avoiders did not emerge in the stressed female group. However, there were poor avoiders in the stressed male group and both control groups. All data are shown as mean ± SEM.

To assess whether the facilitation of SAA in stressed female rats was driven by the reduction in poor avoiders, we analyzed SAA acquisition in only those rats who successfully acquired the AR (i.e., poor avoiders were eliminated from the analysis). As shown in Figure 3A (Left), stressed female rats exhibited facilitated acquisition of ARs relative to controls. This impression was confirmed in an ANOVA that revealed significant main effects of Treatment [F(1, 26) = 6.86, p = 0.01] and Day [F(3, 78) = 58.74, p < 0.0001], as well as a significant interaction between Day and Treatment [F(3, 78) = 4.09, p = 0.0095]. In contrast, stress and control female rats showed similar acquisition of ITRs during training (main effect of Day [F(3, 78) = 8.69, p < 0.0001]) (Figure 3A, Right). Unlike females, stress did not affect acquisition of ARs in males [F(1, 40) = 0.005, p = 0.9418) (Figure 3C, Left); both groups acquired ARs (main effect of Day, [F(3, 120) = 82.64, p < 0.0001]) and ITRs (main effect of Day, [F(3, 120) = 11.55, p <0.0001] similarly (Figure 3C). Thus, stress not only eliminated poor avoiders among females, but it also facilitated the acquisition of the avoidance response.

**Figure 3.**
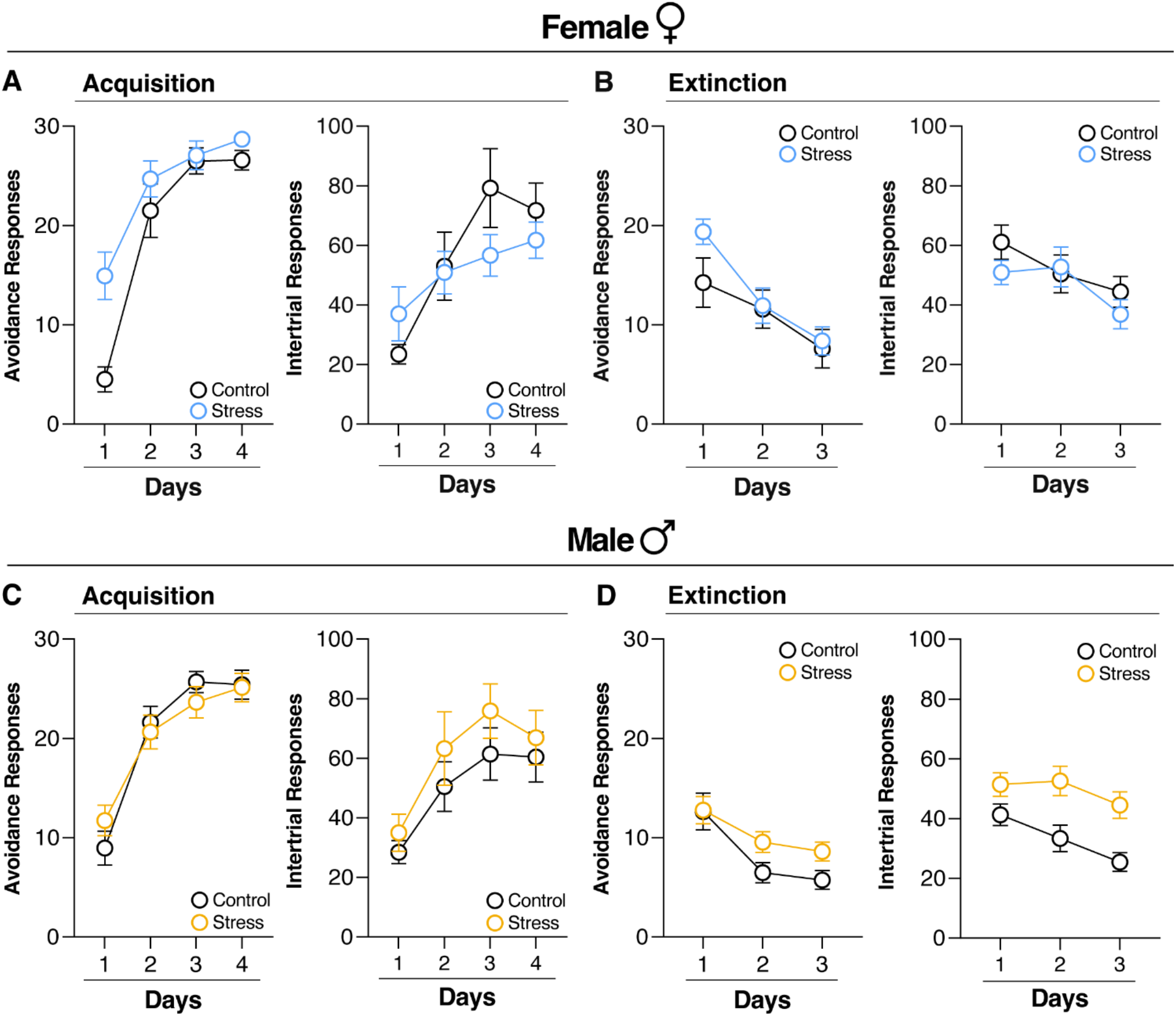
Acute stress facilitates two-way signaled active avoidance acquisition in female rats and induces resistance to extinction in male rats. (A) and (C) (Left) Average number of ARs and (Right) ITRs performed on each day of SAA training excluding poor avoiders. (A) Female rats exposed to acute stress (n = 16) performed significantly higher ARs (Left) during SAA training than their control counterparts (n = 12), but not ITRs (Right). (C) Male rats exposed to acute stress (n = 23) performed similar levels of ARs (Left) and ITRs (Right) to their control counterparts (n = 19) during SAA training. (B) and (D) (Left) Capped ARs and (Right) total number of ITRs performed each day of SAA extinction excluding poor avoiders. (B) Female rats submitted to acute stress performed similar ARs and ITRs each day of SAA extinction. (D) Male rats submitted to acute stress performed significantly higher ARs (Left) and ITRs (Right) than their control counterparts, demonstrating a stress-induced resistance to SAA extinction. All data are shown as mean ± SEM.

To compare male and female rats directly, we focused on the first day of SAA acquisition in which the difference among the groups was most prominent. A two-way ANOVA with the between-subject factors of Treatment and Sex (Male or Female) was performed on ARs from Day 1 of SAA training in all subjects. This analysis revealed a main effect of Treatment [F(1, 66) = 12.63, p = 0.0007] and a significant Sex by Treatment interaction [F(1,66) = 4.21, p = 0.04]. Thus, our results indicate a sex-specific enhancement of SAA acquisition by acute stress, whereby stressed females demonstrated accelerated avoidance learning relative to other groups.

After training, rats were submitted to three days of SAA extinction in either the training context or a novel context (Figure 1A). As shown in Figure 3B and 3D, acute stress induces a resistance to SAA extinction in male but not female rats. To assess the effect of acute stress on ARs and to compare males and females directly, a three-way ANOVA with two between-subjects factors of Treatment (Stress or Control) and Sex and a within-subjects factor of Day was performed on the data. This ANOVA revealed a significant Day x Treatment x Sex interaction [F(2,132) = 4.13, p = 0.0182), confirming the impression that acute stress induces a resistance to AR extinction in male but not female rats (Figure 3B and 3D, left). The same analysis was performed on ITRs (Figure 3B and 3D, right) revealing a significant interaction between Treatment and Sex [F(1,66) = 5.61 p = 0.02], demonstrating that stress induced a resistance to extinction of ARs and ITRs in males, but not females.

Importantly, shuttling during the 5-minute acclimation period was analyzed for impacts of acute stress on locomotion. These data were analyzed using the three-factor ANOVA described above. Importantly, this ANOVA did not reveal a significant main effect of Treatment [F(1,66) = 0.717, p = 0.40), indicating that acute stress did not alter baseline levels of locomotion, and that increased avoidance and intertrial responses can be attributed to a stress-induced increase in CS elicited shuttling.

### Experiment 2: Effect of acute stress on contextual fear conditioning (SEFL)

One explanation for the sex-dependent effects of stress on SAA is a differential effect of stress on Pavlovian freezing that interacts with avoidance learning (Moscarello & LeDoux, 2013; Moscarello & Hartley, 2017). Though previous reports have not found sex differences in the acquisition of fear conditioning following acute stress (Poulos et al., 2015, Hassien et al., 2020; Gonzalez et al., 2021), we performed an experiment to ascertain whether, in our own hands, sex mediates the effects of acute stress on contextual fear conditioning. Male and female rats were submitted to the acute stress procedure, followed a week later by one-trial contextual fear conditioning and retrieval in a distinct context (Figure 1B). As shown in Figure 4A, stressed animals had a significant increase in postshock freezing relative to their controls. This increase in freezing was observed again during retrieval testing in the one-trial conditioning context (Figure 4C), indicating the presence of the SEFL phenotype. To examine the increase in freezing during contextual fear conditioning, a three-factor ANOVA with one within-subject factor of Period (Pre-Shock and Post-Shock) and two between-subjects factors of Treatment and Sex was employed, revealing a main effect of Period [F(1,36) = 23.55, p < 0.0001] and Treatment [F(1, 36) = 27.46, p < 0.0001], as well as a significant interaction between Treatment and Period [F(1,36) = 4.76, p = 0.0357], supporting the impression that stressed animals had a greater increase in post-shock freezing than controls (Figure 4A).

**Figure 4.**
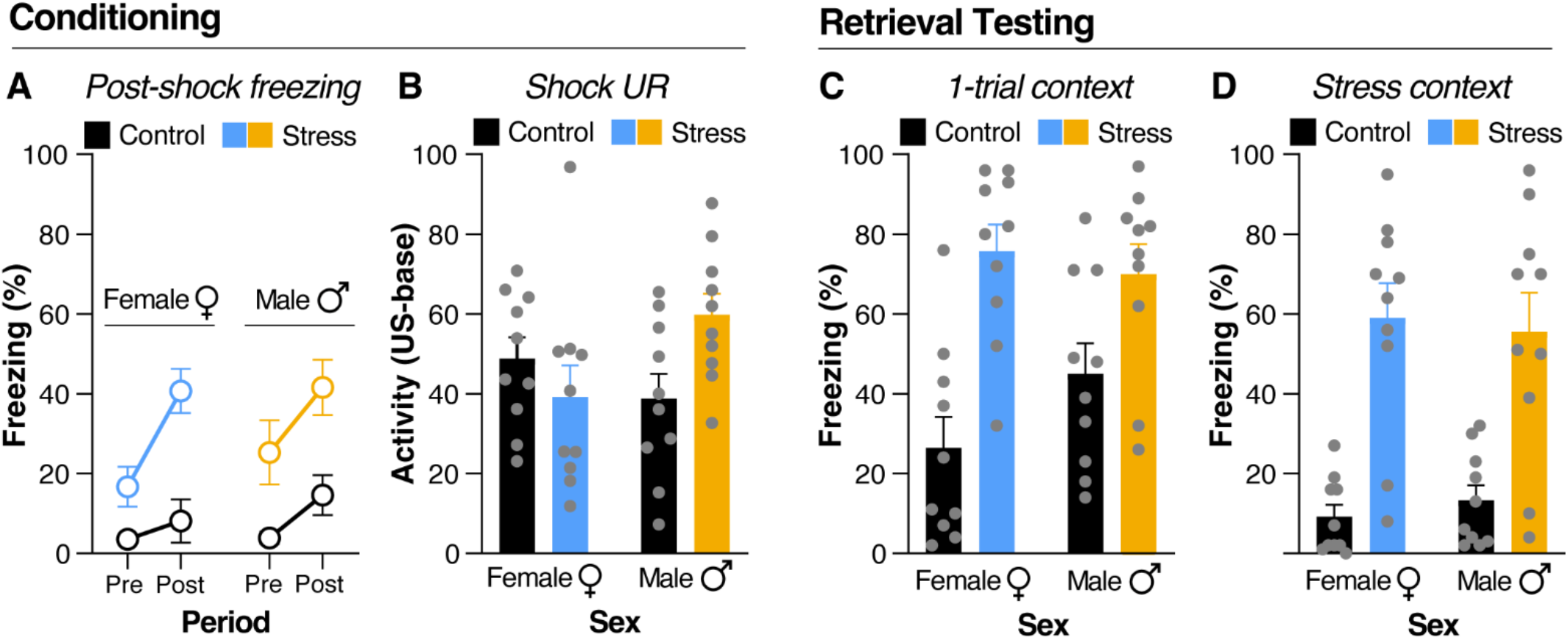
Acute stress induces SEFL in both male and female rats. (A) Freezing in the 3-minute period before and 30 seconds after shock presentation during weak fear conditioning. Male (n = 10) and female (n = 10) rats submitted to acute stress show a significant increase in post-shock freezing during fear conditioning compared to their male (n = 10) and female (n = 10) control counterparts. (B) Unconditioned response (UR) to shock. UR was computed as a difference between load cell activity during the 1-s shock and load cell activity during the 10-s period before the shock. Female rats submitted to acute stress show a reduced UR to shock compared to their control counterparts. Male rats submitted to acute stress show a significant increase in UR to shock compared to their control counterparts. (C) Freezing during the fear retrieval test. Both male and female rats submitted to acute stress show a significant increase in freezing during the context fear retrieval test, indicating the presence of SEFL. (D) Freezing during the stress context retrieval test. Male and female rats submitted to acute stress show significantly higher freezing to the stress context than their control counterparts. All data are shown as mean ± SEM.

To determine whether SEFL was driven by a stress-induced increase in shock reactivity, we examined shock-induced URs during the weak contextual conditioning procedure. The shock UR was computed as a difference between load cell activity (an index of motor activity) during the 1-s shock and load cell activity during the 10-s period before the shock. As shown in Figure 4B, stress sensitized the shock UR in males, but slightly decreased the UR in females. An ANOVA with two between-subjects factors of Treatment and Sex revealed a significant Treatment by Sex interaction [F(1, 36) = 6.059, p = 0.0188] (Figure 4B). Although this sex difference in shock reactivity did not impact SEFL, it is possible that it contributed to facilitated SAA in female rats (though we are not able to quantify shock reactivity in the SAA chambers).

One day after one-trial contextual fear conditioning, animals were submitted to a retrieval test in the same context. As shown in Figure 4C, animals submitted to the acute stress procedure exhibit a significant increase in contextual freezing at retrieval, demonstrating the presence of SEFL. To confirm this, freezing behavior from this test was analyzed by a two-way ANOVA with two between-subjects factors of Treatment and Sex. Results showed a main effect of Treatment [F(1, 36) = 25.18, p < 0.0001], indicating that both male and female animals submitted to the acute stress protocol were demonstrating the same degree of SEFL (Figure 4C).

The following day, rats were placed back into the acute stress context for a retrieval test. Freezing was significantly higher in the stressed animals in comparison to the control animals. An ANOVA identical to that performed on the contextual fear retrieval data revealed a main effect of Treatment [F(1, 36) = 43.29, p < 0.0001], indicating that stressed animals froze more than controls regardless of sex (Figure 4D). Taken together, these data reveal similar effects of inescapable shock on freezing behavior in male and female rats. This indicates that the effect of acute stress produces a sex-dependent facilitation of aversive learning that is selective for the acquisition and extinction of SAA.

## DISCUSSION

Here, we present data demonstrating that acute stress has a sex-dependent effect on the acquisition and extinction of SAA. In females, acute stress facilitated the acquisition of SAA by increasing ARs on the first day of SAA training and eliminating poor avoiders. Despite enhanced acquisition of avoidance, females that received acute stress displayed normal SAA extinction. Conversely, males submitted to the same acute stressor acquired SAA like control animals but demonstrated increased ARs and ITRs during extinction. Finally, consistent with previous reports, we did not observe a sex difference in the effects of SEFL on contextual freezing (Poulos et al., 2015, Hassien et al., 2020; Gonzalez et al., 2021). Thus, we can conclude that acute stress facilitated SAA in a sex-dependent manner that is specific to avoidance and is not related to changes in defensive freezing or Pavlovian fear conditioning.

Sex differences have been observed in both PTSD symptoms and aversive learning processes, and these differences are thought to underlie PTSD (Maren et al., 1994; Kessler et al., 1995; Fullerton et al., 2001; Gupta et al., 2001; King et al., 2013; Hourani et al., 2015; Panayiotou et al., 2017; Sheynin et al., 2017; Merz et al., 2018; Wen et al., 2022). Despite these differences, very few studies examining the impact of stress on avoidance consider sex as a biological variable (Dalla et al., 2007; Shors et al., 2007, McDowell et al., 2015, Collins et al., 2023). Of those that do, results do not provide a translationally valid model for sex differences in PTSD. For example, Knox and colleagues (2023) observed increased unconditioned avoidance in stressed female rats in a light/dark avoidance task after single prolonged stress (SPS), a procedure that models features of PTSD (Liberzon et al., 1997; Knox et al., 2012; Souza et al., 2017). However, the canonical SPS-induced extinction retrieval deficit (Knox et al., 2012) was absent in both male and female rats, making it difficult to interpret these results. In the learned helplessness paradigm, males submitted to uncontrollable shock demonstrate an impairment in the acquisition of a shuttle-box escape task, whereas males that were able to control shock acquire the escape contingency (Maier et al., 1979). However, females show a resistance to uncontrollable shock and do not show learned helplessness in shuttle box escape (Dalla et al., 2007; Shors et al., 2007). Conversely, chronic restraint stress does not impact lever press avoidance performance in males but impairs lever press avoidance in females (McDowell et al., 2015). These stress-induced impairments may be more akin to depressive-like phenotypes and highlight the need to model behavioral disturbances in both sexes to conduct translationally valid, preclinical research on PTSD and anxiety.

The sex-dependent pattern of results reported here suggests a clinically relevant impact of acute stress on avoidance. In the SAA paradigm, females submitted to the acute stress procedure rapidly acquire the avoidance contingency, demonstrating a sex-specific stress-induced sensitization of avoidance learning that mirrors the increased prevalence of avoidant symptoms in women with PTSD (Kessler et al., 1995; Fullerton et al., 2001; Panayiotou et al., 2017). In contrast, the stress-induced impairment in SAA extinction observed in male rats may model hypervigilant behavior, which is more prevalent in men with PTSD (King et al., 2013; Hourani et al., 2015). Both avoidance and hypervigilance are highly predictive of posttraumatic functional impairment (Norman et al., 2007), and early expression of hypervigilance after trauma predicts minimal overall symptom improvement (Schell et al., 2004). Thus, the data presented here constitute a valuable model for investigating the sex-specific symptoms of PTSD that are critical for the maintenance of the disorder.

Though the neural substrates mediating the differential influence of stress on SAA are unknown, prior research suggests some candidate regions. One possibility is that acute stress exerts its effects via the infralimbic cortex (IL), a medial prefrontal cortical region that is critical for instrumental avoidance learning (Moscarello and LeDoux, 2013; Capuzzo and Floresco, 2020) and the extinction of conditioned fear (Giustino and Maren, 2015; Quirk and Mueller, 2007; Milad and Quirk, 2002). For example, inactivation of the IL undermines SAA acquisition (Moscarello and LeDoux, 2013). Importantly, the effects of stress on IL are sex-dependent (Farrell et al., 2014; Binette et al., 2023). Several reports demonstrate that stress has dramatic effects on neuronal morphology in IL (Izquierdo et al., 2006; Lehmann and Herkenham, 2011; Moench et al., 2016) and impairs IL function in males (Pennington et al., 2017; Canto De Souza, 2021; Nawreen et al., 2020). However, it has also been shown that stress facilitates performance in IL-dependent tasks in female rats (Bowman et al., 2001, 2003; Beck and Luine, 2002; Conrad et al., 2003; Maeng and Shors, 2013, Garret and Wellman, 2009; Yuen et al., 2016). Thus, facilitated SAA acquisition and normal SAA extinction in females, along with impaired extinction in males, are broadly consistent with divergent, sex-specific effects of acute stress on IL, whereby IL function is somewhat impaired in males and facilitated in females.

The bed nucleus of the stria terminalis (BNST) is another candidate region that may mediate the behavioral effects of stress on avoidance. The BNST receives convergent projections from multiple regions implicated in SAA, such as IL (Moscarello & LeDoux, 2013; Glover et al., 2020), central amygdala (Moscarello & LeDoux, 2013; Asok et al., 2018), and ventral hippocampus (Lee and Davis, 1997; Oleksiak et al., 2021). In addition, the BNST regulates the impact of stress on aversive learning processes in males, but not females (Bangasser and Shors, 2010), suggesting that it may contribute to stress effects on extinction in male rats reported here. Indeed, chemogenetic activation of BNST in male subjects facilitates the expression of ARs in SAA (Guerra et al., 2023). The BNST is also sexually dimorphic (Allen and Gorski, 1990; Shah et al., 2004; Uchida et al., 2019; Tsukahara and Morishita, 2020), and the action of gonadal steroids and their metabolites in the BNST may play a role in sex-dependent aversive learning processes, such as contextual fear conditioning (Maren et al., 1994; Nagaya et al., 2015; Blair et al., 2022). Because acute stress enhanced the expression of ARs and ITRs in males under extinction conditions, we propose that stress induced BNST plasticity may explain the resistance to SAA extinction observed in stressed males.

In summary, the data presented here demonstrate sex-specific impacts of acute stress on avoidance in SAA, whereby stressed females display enhanced acquisition of SAA and stressed males display a resistance to SAA extinction. Our results add to the literature assessing the impact of stress on avoidance by including sex as a biological variable (Shors et al., 2007; Dalla et al., 2008; McDowell et al., 2015; Collins et al., 2023) and providing a translationally relevant model for avoidance in PTSD in both males and females. Further work will explore the mechanism behind the sex differences observed here, with the goal of providing sex-specific treatments for PTSD.

## Author contributions

S.P., J.M., and S.M. designed all the experiments and analyses. S.P., C.O., C.P., and C.M. performed behavioral experiments. S.P. analyzed the data. S.P., J.M., and S.M. wrote the manuscript.

